# The photocycle of orange carotenoid protein conceals distinct intermediates and asynchronous changes in the carotenoid and protein components

**DOI:** 10.1101/167478

**Authors:** E.G. Maksimov, N.N. Sluchanko, Y.B. Slonimskiy, A.V. Stepanov, E.A. Shirshin, G.V. Tsoraev, K.E. Klementiev, O.V. Slatinskaya, E.P. Lukashev, T. Friedrich, V.Z. Paschenko, A.B. Rubin

## Abstract

The 35 kDa water-soluble Orange Carotenoid Protein (OCP) is responsible for photoprotection in cyanobacteria. It acts as a light intensity sensor that simultaneously serves as efficient quencher of phycobilisome excitation energy as well as of reactive oxygen species. Photoactivation triggers large-scale conformational rearrangements to convert OCP from the orange OCP^O^ state to the red active signaling state OCP^R^, as demonstrated by various structural methods. Eventually, such rearrangements imply complete yet reversible separation of structural domains (C- and N-terminal domain) and significant translocation of the carotenoid cofactor. Very recently, dynamic crystallography of OCP^O^ crystals suggested the existence of photocycle intermediates with small-scale rearrangements that may trigger further transitions in the protein. However, the currently existing gap between the ultra-fast picosecond and 100 millisecond time scale of spectroscopic and structural data precludes knowledge about distinct intermediate states. In this study, we took advantage of single 7 ns laser pulses to study carotenoid absorption transients in OCP on the time-scale from 100 ns to 10 s, which allowed us to detect a red intermediate state preceding the red signaling state OCP^R^. In addition, time-resolved fluorescence spectroscopy and following assignment of carotenoid-induced quenching of different tryptophan residues revealed a novel orange intermediate state, which appears during back-relaxation of photoactivated OCP^R^ to OCP^O^. Our results show asynchronous changes in the carotenoid and protein components and provide refined mechanistic information about the OCP photocycle as well as introduce new kinetic signatures for future studies of OCP photoactivity and photoprotection.

**Significance statement:** Cyanobacteria utilize the Orange Carotenoid Protein (OCP) to protect their photosynthetic apparatus from the harmful effects of intense sunlight. OCP is a blue light-triggered photoswitch, which undergoes photoconversion from its dark adapted orange to the active red state, the latter being able to interact with the phycobilisome antennae and quench their fluorescence, thus avoiding excessive energy flow to the photosystems. With the help of the fluorescence recovery protein (FRP), OCP detaches from phycobilisomes and can return faster into the orange state. Until now, only the thermodynamically stable orange state and the metastable red state are established in a primitive photocycle. In this work, we apply transient absorption and fluorescence spectroscopy and identify two novel photocycle intermediates of physiological relevance.

## 1. Introduction

Adapted to perceive the macroworld, human senses constantly fail to acknowledge the most incredible yet ordinary capabilities of biological materials on the nanoscale. Translated into units we can oversee, the absorption of a single green 500-nm photon by a 35-kDa protein compares to the traceless swallowing of a 4-g, 9-mm Parabellum bullet, fired from a shotgun at the speed of sound, by a 35-g slab of soft biological matter. This apocalyptic event, taking place in myriads at every twinkle of the eye, forms the basis of oxygenic photosynthesis for the benefit of Life on the Planet. And it is also the task of the 35-kDa Orange Carotenoid Protein (OCP) in cyanobacterial photoprotection, which utilizes the extraordinary capability of its carotenoid cofactor to convert electronic excitation energy by internal conversion within a few picoseconds into harmless heat in order to shield the sensitive photosystems from excessive excitation energy arriving at the phycobilisome (PBs) antennae complexes at high light intensities (1-9). OCP is a blue-green light photoreceptor capable of undergoing a red shift of its absorption spectrum upon illumination (10, 11). Photoactivity necessarily requires a single 4-ketolated carotenoid molecule to be coordinated by OCP, which is 3’-hydroxyechinenone when purified from cyanobacteria (3, 12), but also incorporation of echinenone (4-keto; ECN) and the 4,4’-diketolated canthaxanthin (CAN) preserves photoactivity, whereas binding of the 3,3’-dihydroxylated zeaxanthin precludes spectral switching (13-15). OCP is divided into two domains of about half its molecular weight, with an all-α-helical N-terminal domain (NTD) and a mixed α/β-type C-terminal domain (CTD) bearing similarity to the protein family of nuclear transfer factors (NTF-2, Pfam: PF02136), which almost symmetrically enclose a central carotenoid-binding cavity that nearly completely shields the cofactor from the aqueous environment (16, 17). The orange state OCP^O^ is structurally stabilized by distinct carotenoid-protein interactions including specific H-bonds from Trp-288 and Tyr-201 in the CTD to the 4-keto oxygen at one of the β-rings, as well as interactions to Tyr-44 and π-stacking with Trp-110 in the NTD, but also extensive protein-protein interactions involving multiple H-bonds across the extensive NTD-CTD inter-domain interface (including two formed between Asn-104 and Trp-277, and Arg-155 and Glu-244 (18)) and binding of the αA-helix of the L-shaped N-terminal extension (NTE) to the central β-sheet surface on the CTD (19-22).

Upon photoconversion from the orange OCP^O^ state, the red-shifted OCP^R^ state is formed, which can interact with phycobilisome antennae complexes and quenches their fluorescence in the photoprotective mechanism of non-photochemical quenching (7, 21, 23, 24). According to biochemical (size-exclusion chromatography, SEC) and structural (small-angle X-ray scattering, SAXS, or static and dynamic X-ray crystallography) studies, the active, quenching OCP^R^ state is characterized by increased disorder in terms of a molten globule, detachment and unfolding of the αA-helix from the CTD, domain separation and, eventually, a carotenoid translocation by 12-Å into the NTD, thereby forming a spectral state indistinguishable from the one of the Red Carotenoid Protein (RCP). The latter is a hydrolysis product of OCP comprising only the carotenoid-bound N-terminal domain, which constitutively quenches PBs fluorescence (25-28). The photoprotective mechanism in *Synechocystis* is terminated by the action of the Fluorescence Recovery Protein (FRP), which promotes dissociation of OCP^R^ from the PBs and accelerates recovery to the OCP^O^ state (29-31). Binding of FRP occurs to the CTD, most likely at a site covered in OCP^O^ by the αA-helix, although some secondary, as yet unidentified, site might necessarily exist on the NTD or the interdomain linker as well to enable reversion of domain separation and back-sliding of the carotenoid cofactor into its initial position (22, 32). Spectral and structural changes of the protein and the carotenoid upon photoexcitation have been studied by multiple spectroscopic approaches (14, 28, 33, 34); however, main spectral and structural intermediates within the OCP photocycle (if existent) are still elusive, as is the case for the atomic structure of the most important, biologically active OCP^R^ state.

The photocycle of OCP was previously studied with a time resolution of up to 100 ms by many scientific groups including ours. Such experiments are usually used for determining the rate of the OCP^R^ to OCP^O^ transition, which is often considered as the rate of protein refolding. Femtosecond time-resolved absorption spectroscopy has been used to probe the initial events following photoexcitation and excited state dynamics covering the time scale up to nanoseconds, which have revealed short-lived far-red-shifted absorption spectra, which have frequently been discussed as bearing similarities to internal charge transfer (ICT) states (2, 34, 35). The ultra-fast (fs) transitions between the excited states of the OCP chromophore were studied for several OCP species revealing that in both, orange and red, states the ECN cofactor has a very short (~ 6 ps) excited state lifetimes indicating that both states are potentially suitable for dissipation of excitation energy of PBs, thus, the conformation of the protein matrix, but not the spectral state of its chromophore, are crucial for PBs fluorescence quenching (2, 35). Additionally, substitution of the Arg-155 to Leu suppresses the induction of fluorescence quenching without the loss of photoactivity (18) from the one side, and constitutively quenching forms like RCP and the OCP-W288A mutant (17, 21) indicate that photoconversion and quenching are discrete events, suggesting that the protein state determines its functionality, while the color of the sample is only a side effect. As the rates of protein motions are significantly slower than the fs transients, the physiologically relevant OCP state capable of PBs fluorescence quenching should appear when the carotenoid is already back in the ground state, while between 100 ms and 1 s, there is already a decay of the red state due to back-relaxation and refolding of the protein.

Raman spectroscopy and theoretical calculations have hinted at the configurational changes of the carotenoid that give rise to spectral shifts, and at the influence of the protein matrix on color tuning (11, 15, 33). Photoconversion is apparently accompanied by a reduction of the ring rotation angles at the C6-C7(C6’-C7’) single bonds of the carotenoid, which leads to an increase of the apparent conjugation length of the polyene chain, and to a relief of bending tension of the latter. As outlined by the recent study of Bandara and colleagues (36), this strained chromophore configuration in the dark-adapted state distinguishes OCP from other photoreceptors, in which strained chromophores are the initial or intermediate photoproducts on the way to the product state. Bending of the chromophore in OCP^O^ most likely is due to the off-axis H-bond interaction of the 4-keto oxygen to the indole N-H group of Trp-288 in the CTD, while the carotenoid is otherwise tightly packed by a cavity constriction at the inter-domain interface and mainly hydrophobic and π-stacking interactions with Trp-110 in the NTD. Dynamic X-ray crystallography has identified very early structural intermediates of photoconversion showing detachment of the αA-helix, initial rigid-body separation of the CTD and retraction of Tyr-201 and Trp-288 from the terminal β-ring, indicating that breaking of the specific chromophore-protein H-bonds is one of the initial triggers of photoconversion (36). At the other end of the time scale, in the range of hundreds of milliseconds to hundreds of seconds, spectroscopic monitoring of light-induced absorption or fluorescence changes has revealed the kinetic and thermodynamic aspects of the global phototransformations between OCP^O^ and OCPR (11), as well as the influence of amino acid substitutions, or the action of FRP on these processes (20-22). However, there is still a large gap of time scales to be covered by time-resolved absorption and other spectroscopic techniques in order to elucidate processes occurring between hundreds of nanoseconds to milliseconds for resolving the intermediate states of OCP, which appear during the photocycle. This previously omitted time window spans many orders of magnitude after photoexcitation, when the chromophore has for long returned to the electronic ground state, preceding the dramatic conformational changes of the protein, which occur upon its activation and relaxation. Importantly, this range of timescales (10^-7^-10^-3^ s) has been demonstrated to be crucial for the photocycle of other photoactive proteins (37-39), covering several transitions between the intermediate states.

In this paper, we made use of the flash photolysis and picosecond fluorescence spectroscopy techniques to reveal novel OCP red and orange intermediate states characterized by distinct spectroscopic features, which appear during phototransformation to the OCP^R^ product state and back-relaxation to the dark-adapted OCP^O^ state. These findings result in a more comprehensive and complete photocycle model of OCP with implications on its biological function and regulatory interactions relevant for the role of the protein *in vivo*.

## 2. Results and Discussion

### 2.1. Single flash-induced absorption transients of OCP reveal red intermediate state

First of all, our transient absorption experiments (*Figure 1*) show that after irradiation of the sample with a 7-ns laser flash, the increase of OCP absorption at 550 nm (i.e. its red shift) occurs faster than 200 ns, which is the time resolution of our instrument. This remarkable feature indicates that only small changes of chromophore–protein interactions are necessary for the spectral conversion of OCP^O^ to a state with red-shifted chromophore absorption. Namely (? This is because), 200 ns after photon absorption, the spectral characteristics of OCP are already similar to those of OCP^R^, but the protein state could yet be significantly different from the physiologically active state (i.e. capable of PBs fluorescence quenching). This is based on the assumptions that (1) conformational rearrangements of OCP are necessary for the interaction with the binding site on PBs [16-17, 25-28] and (2) no dramatic structural changes of the protein could be exerted on a <200 ns time scale.

Importantly, the amplitude of the absorption changes saturated at pulse energies above 30 mJ (*Figure 1B*), indicating that all carotenoid molecules were excited. This fact allows calculating the quantum yield of OCP photoconversion to be equal to 0.3 % and the yield of OCP^R^ formation ~ 0.2 %, in excellent agreement with our previous estimations (11). Such a low yield of photoconversion is probably related to the fact that activation of PBs quenching is achieved efficiently only at high intensity of solar radiation. If the yield would be higher, OCP photoconversion leading to PBs quenching may occur already unter potentially harmless conditions, thus diminishing the efficiency of light harvesting. On the other hand, lower photoconversion yield would have reduced the efficiency of photoprotection, so the proper balance between the stability of OCP^O^ and its sensitivity to blue-green light is crucial.

Further changes of the OCP absorption at 550 nm can be described by three exponential decay components with significantly different lifetimes ranging from hundreds of μs to tens of s (at 36 °C). The time constant of the first (fast, C1) component is equal to 300 μs, the second (C2) is about 18 ms and the third (C3) one is about 3.3 s (*Table 1*). It should be noted that the characteristic rate constant of the slow C3 component has the same order of magnitude as the R-O back-conversion rate, which was determined with a 100 ms resolution technique (~ (9 s)^-1^) at the same temperature (11, 21).

In order to further investigate the origin of the two faster C1 and C2 components in the transient absorption decay, we tested several factors affecting the rates of photoconversion, such as the effect of FRP, temperature and high concentrations of phosphate, with the latter expected to act as a kosmotrope and stabilize the OCP^O^ state (40). Addition of FRP slightly increased the rate of C1 to (220 μs)^-1^, the C2 component vanished, and the rate of C3 was significantly increased up to ~ (95 ms)^-1^. Importantly, the increase of the rates in the presence of FRP enabled us to reduce the integration time to solve the principal problem of such time-resolved experiments requiring the accumulation of hundreds and thousands of kinetics traces, i.e., several days of continuous measurements. Addition of FRP also made it possible to accurately measure the temperature dependency (*Figure 1* and *Supplementary Figure 1*) of these rates and to estimate the corresponding activation energies, which appeared to be almost identical for the C1 and C3 components (~ 22 kcal/mol, see *Figure 1C*). The similarity of the activation energies indicates that the huge differences in the rates of C1 and C3 are due to the pre-exponential factor in the Arrhenius law, which reflects the probability of product formation in each of the elementary steps. Thus, the fastest C1 and the slowest C3 components reflect two different states with red shifted absorption spectrum, for which relaxation to the orange state is characterized by different numbers of attempts (or “collisions” in collision theory) necessary to restore spectral characteristics of OCP^O^. One may assume that this is due to the protein conformation of OCP at different stages of photoconversion: protein conformation of the fast-decaying red intermediate (further OCP^RI^) is probably very similar to that of OCP^O^ (small number of attempts to get to initial OCP^O^), while the slow decay is a feature of the commonly recognized OCP^R^ with separated structural domains (very high number of attempts to get OCP^O^). Assuming that domain separation and 12-Å carotenoid translocation into the NTD correctly describe the physiologically active OCP^R^ state, as inferred from crystal structures (26) and resonance energy transfer measurements (28), the slow component reflects the fact that the carotenoid needs to penetrate into the CTD again in order to recover its orange absorption spectrum. Thus, spatial separation of CTD and NTD significantly reduces the probability of back-conversion to the stable OCP^O^ state.

**Table 1.** Results of approximation of flash-induced absorption transients of OCP by multiexponential decay shown in *Figure 1A*. All experiments were conducted at 36 °C. The FRP to OCP (or ΔNTE OCP) concentration ratio was equal to 1.6.

| Sample | Fast component (C1) |  | Middle component (C2) |  | Slow component (C3) |  |
| --- | --- | --- | --- | --- | --- | --- |
|  | t<sub>1</sub>, μs | a<sub>1</sub>, % | t<sub>2</sub>, ms | a<sub>2</sub>, % | t<sub>3</sub>, ms | a<sub>3</sub>, % |
| OCP | 300±20 | 31 | 18±3 | 12 | 3300±100 | 57 |
| OCP in 1M NaPi | 295±20 | 59 | - | - | 430±50 | 41 |
| OCP + FRP | 220±20 | 32 | - | - | 95±5 | 68 |
| ΔNTE OCP + FRP | 70.4±4 | 24 | - | - | 272±20 | 76 |

In order to test how the separation of domains affects the yields and lifetimes of different OCP intermediates with red-shifted absorption, we used the non-specific action of high concentrations of inorganic phosphate, a well-known kosmotrope, which is always present in buffers for experiments with PBs, thereby affecting the results of every *in vitro* fluorescence quenching experiment (16, 24). We found that phosphate significantly reduces the amplitude of flash-induced spectral changes at 550 nm, accelerating the slow С3 component (*Table 1*) and simultaneously reducing its yield (*Figure 1*). This observation indicates that high concentrations of phosphate likely hinder domain separation of OCP, which is manifested in the rate of its back-conversion, and, therefore, accumulation of the physiologically active OCP^R^ state, which explains why *in vitro* PBs quenching occurs only at a huge excess of OCP (~ 40 – 100 OCP per 1 PBs (16, 21, 23, 24)).

At the same time, addition of phosphate does not affect the C1 component rate, thus confirming that the corresponding short-lived OCP^RI^ intermediate, in terms of protein structure, still resembles the compact and stable OCP^O^, and its formation does not require significant conformational rearrangements and occurs before them. In contrast to the effect of phosphate, the rate of C1 is slightly increased by addition of FRP. This effect is relatively small (~ 10 %) compared to the influence of FRP on the C3 component (*Table 1),* indicating that FRP interacts effectively with the physiologically relevant OCP^R^ state, but interactions with the OCP^O^-like OCP^RI^ state are also possible. This conclusion is in agreement with our previous results about the OCP interaction with FRP (21). Next, deletion of the OCP^O^ stabilizing N-terminal extension (NTE), which was recently suggested to also cover the FRP binding site on the CTD of OCP^O^, results in formation of stable FRP-OCP complex even in the dark (22). In such complexes, the rate of the fast C1 component is at least four times faster compared to full-length OCP and three times faster compared to full-length OCP in presence of FRP (*Table 1*). Thus, FRP increases the probability of orange form restoration not only from the physiologically active OCP^R^ state, but also from the intermediate red OCP^RI^ state. These facts might indicate that in the intermediate OCP^RI^, the αA helix of the NTE could already detach from the CTD uncovering the FRP binding site (22). A possible explanation of this phenomenon could be slow detachment of FRP from OCP, which occurs slower than the changes of carotenoid absorption on the timescale exceeding intervals between the excitation laser pulses in our experiment – 1 s, or due to FRP binding to OCP in the orange state, which we observed at high concentrations (21, 32). Although we support the idea of a secondary FRP binding site in the NTD of OCP (32), it seems reasonable to assume that the mechanism of FRP action is not limited to trivial coupling of the OCP domains. If this would be the case, we would have expected to see a reduction of the yield of the slow component for ΔNTE OCP in the presence of FRP, as observed in experiments with phosphate; however, our data indicate that strong binding of FRP to ΔNTE OCP does not prohibit formation of the OCP^R^ state with separated domains (*Figure 1*).

**Figure 1.**
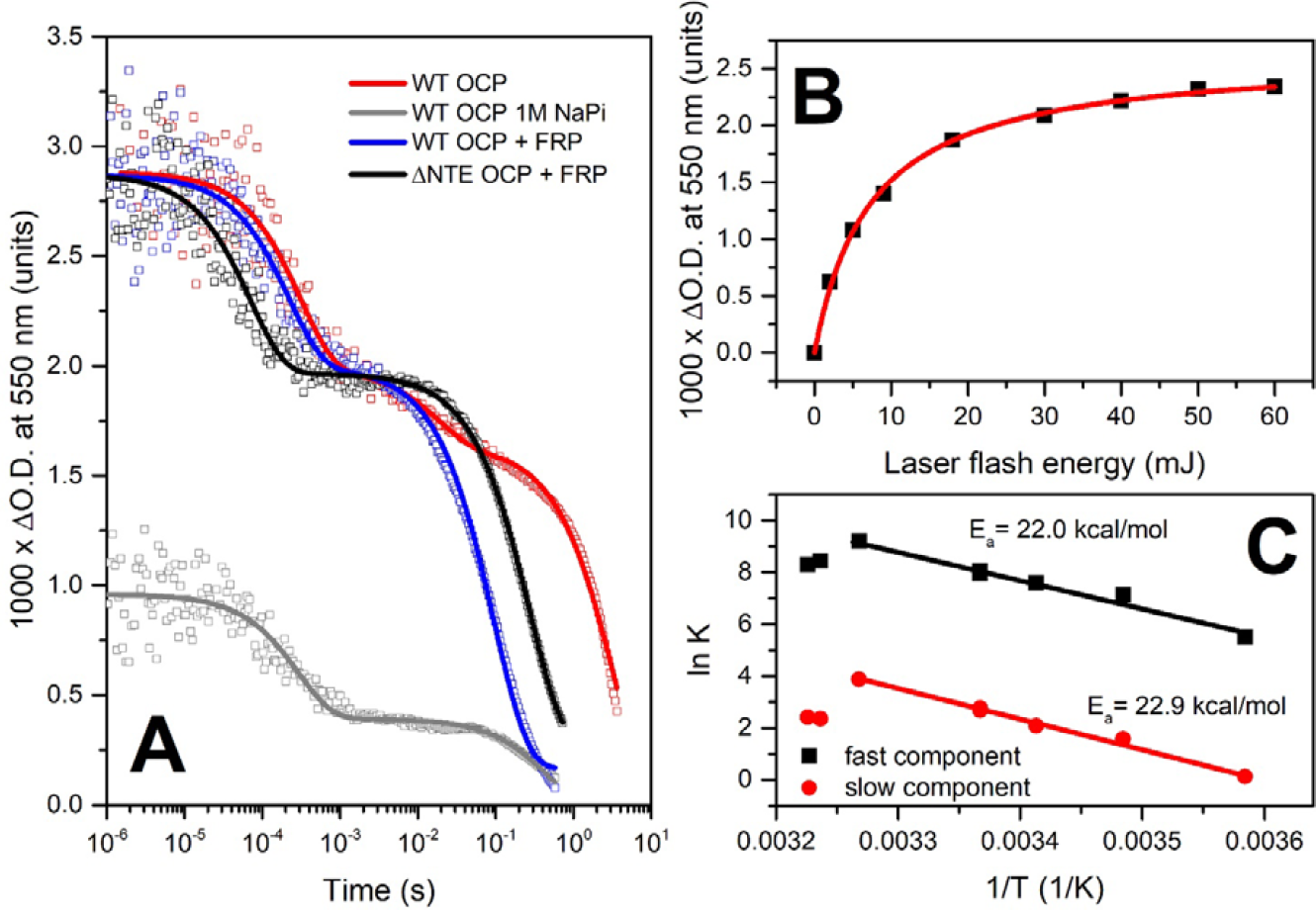
(**A**) – flash-induced transitions of OCP absorption at 550 nm approximated by multiexponential decay: OCP – red, OCP in 1M phosphate – grey; OCP in the presence of FRP (1/1.6 concentration ratio) – blue; ΔNTE OCP in presence of FRP (1/1.6 concentration ratio) – black. Note the logarithmic timescale covering almost 7 orders of magnitude. (**B**) – dependency of the photoconversion amplitude on the energy of the laser flash for OCP in the presence of FRP (1/1.6 concentration ratio). (**C**) – Arrhenius plot of the fastest (C1) and the slowest (C3) components’ rates of absorption changes of OCP in the presence of FRP. Experiments were conducted in the range of temperatures from 6 to 37 °C.

### 2.2. Intrinsic fluorescence of OCP reveals an orange intermediate state

Considering the lifetimes of OCP^RI^ and OCP^R^ states (~ 300 μs and 3 s, respectively, – see *Table 1*) it is clear that continuous (or high frequency) excitation of the sample by actinic light leads to accumulation of physiologically active OCP^R^ state. Kinetics of back-relaxation of this state to OCP^O^ was previously studied in detail, but from the perspective of the chromophore absorption spectrum only. Previously, we showed that the OCP photocycle can be studied by different fluorescence techniques (14, 28, 41), including the monitoring of Trp fluorescence (11). Thus, we decided to follow the intrinsic fluorescence of OCP and compare the kinetics of carotenoid- and protein-related transients. For this purpose, we used the same samples and experimental settings as described above to measure the kinetics of OCP^R^ to OCP^O^ relaxation. Namely, (i) changes of absorption at 550 nm and (ii) Trp fluorescence emission at 350 nm were studied. Fluorescence was detected by time-correlated single photon counting, which allowed us to assess not only the changes of fluorescence intensity, but also the lifetime(s) of Trp fluorescence. After photoconversion of the sample into the OCP^R^ state (1 min of actinic light illumination), actinic light was turned off and a sequence of picosecond-resolved fluorescence decay kinetics was recorded (*Figure 2A*). We found that Trp fluorescence intensity is significantly (up to 85 %) quenched in the dark-adapted OCP^O^ state compared to the situation in OCP^R^. However, the corresponding differences of average fluorescence lifetime were less than 40 % (*Figure 2C*), indicating that in the orange state, some Trp residues (2 of 5) may transfer excitation energy to the carotenoid via Förster resonance energy transfer (FRET), while some residues (other 3 of 5) are quenched due to ground state interactions with the carotenoid (representing static quenching). The residues involved in static quenching are most likely directly involved in protein-chromophore interactions, such as Trp-288 (H-bond to carotenoid) and Trp-110 (π-stacking with carotenoid), as inferred from the crystal structures (see the OCP structure overview in *Figure 4A*).

**Figure 2.**
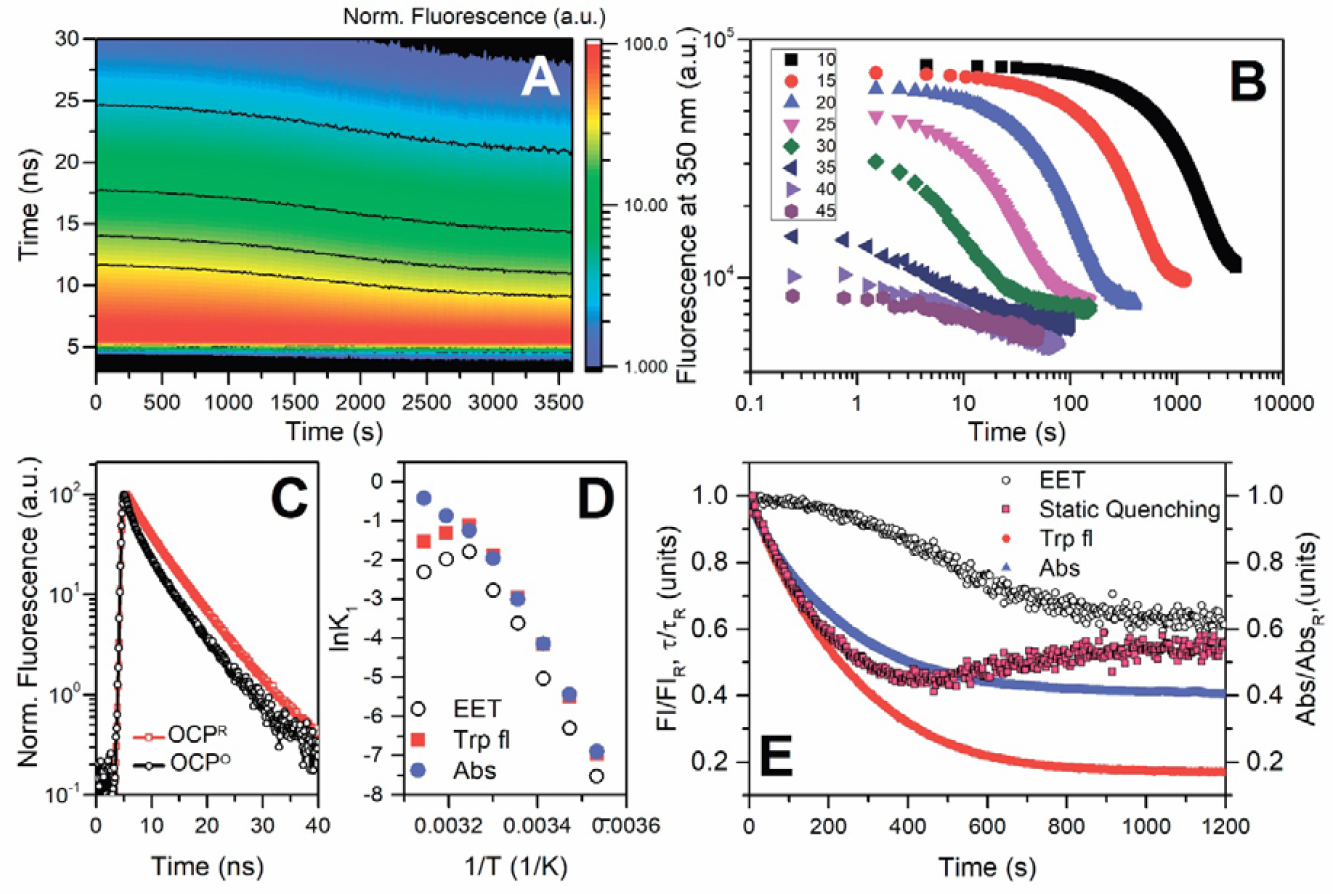
Changes of intrinsic Trp fluorescence of OCP associated with its R-O conversion. (**A**) – a typical set of 400 fluorescence decay kinetics normalized to maximum intensity measured at 10 °C successively after switching off the actinic light, causing conversion of OCP from the red to the orange form. Black lines indicate the levels of fluorescence intensity. (**B**) – time courses of fluorescence intensity changes at 350 nm upon the OCO^R^-OCP^O^ transition at different temperatures. Each experimental point was obtained by integration of the Trp fluorescence decay curve. Note the logarithmic scales of both axes. (**C**) – normalized Trp fluorescence decay kinetics of the photoactivated (OCP^R^) and back-converted (OCP^O^, after 60 minutes in the dark) protein. (**D**) – Arrhenius plots for the OCP^R^-OCP^O^ relaxation measured as the absorption changes at 550 nm (blue circles) and Trp fluorescence changes of intensity (red squares) and average Trp fluorescence lifetime (open circles) calculated from the data presented in (**B**). (**E**) – characteristic time courses of OCP^R^-OCP^O^ transitions of OCP measured as changes of carotenoprotein absorption (blue) at 550 nm, and Trp fluorescence intensity (red circles) and lifetime (or EET, open circles) at 350 nm. Transitions were measured at 15 °C, the concentration of the sample was identical for fluorescence and absorption measurements. Values of O.D., fluorescence intensity and lifetimes were normalized to unity. Time-course of the static quenching (red squares) was calculated from the changes of overall fluorescence intensity 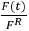 and average lifetimes 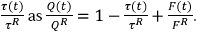

Analysis of the time courses of absorption and fluorescence changes upon back-conversion of the OCP^R^ to OCP^O^ revealed that changes of Trp fluorescence lifetimes occur significantly slower than the overall changes in fluorescence intensity, whereas the rate constants of the latter were similar to those of re-formation of the orange state monitored by absorption changes at 550 nm (*Figure 2D*). This fact indicates that restoration of the initial EET efficiency between Trp residues and the carotenoid, as characteristic for the dark adapted OCP^O^ state, occurs only after the protein-carotenoid system is already in its orange form. On the other hand, the comparison of the changes of fluorescence lifetimes and intensities allows discriminating between dynamic (EET-related) and static quenching (*Figure 2E*). Our data show that restoration of static quenching (e.g. due to H-bond formation or other interactions of Trp residues) precedes the changes in carotenoid-protein absorption. We postulate that formation of this distinct orange intermediate (further termed OCP^OI^) state precedes the final structural rearrangements and restoration of protein-chromophore interactions, which are necessary for stabilization of CTD-NTD interactions and formation of the initial OCP^O^ state. Considering the fact that restoration of the orange absorption spectrum of the carotenoid requires its correct positioning between CTD and NTD of OCP including the formation of H-bonds and eventually also the characteristically bent structure of the polyene chain, we assume that relocation of carotenoid from NTD to CTD is a primary step on the way to the dark adapted stable OCP^O^. As discussed earlier, the rate of OCP^R^-OCP^O^ back-conversion depends on multiple factors; however, the re-establishment of the critical H-bonds between the keto oxygen in the β1-ring of ECN and Trp-288 and Tyr-201 must be crucial for stabilization of OCP^OI^ intermediate state and following formation of OCP^O^.

### 2.3. Roles of Trp-288 and Tyr-201 in protein-carotenoid interactions and stabilization of the orange form

Previously, we characterized the purple W288A mutant of OCP, which appeared to be functionally similar to the physiologically active OCP^R^ state in terms of capability of PBs fluorescence quenching and interaction with FRP (14, 21). Upon addition of FRP to OCP^W288A^ we observed complex formation with 1 to 1 stoichiometry, and the absorption spectrum of this complex exhibited characteristic signatures of the orange state (referred to as “oranging”), presumably, due to OCP^W288A^ stabilization by FRP (21). We hypothesized whether the FRP-induced stabilization of OCP^W288A^ can be mimicked by kosmotropic phosphate anion, which acts non-specifically (19, 24, 40, 42), resulting in stabilization of CTD-NTD interactions (see *Figure 1* and description in text). Increasing the phosphate concentration in OCP^W288A^ solutions (pH was kept constant) caused a gradual increase of absorption in the blue spectral region and the appearance of vibronic substructure, a characteristic signature of the orange OCP form. Surprisingly, at 0.8 M phosphate, the spectrum of OCP-W288A barely retained features of its initial (phosphate-free) state (red-purple, with maximum at 520 nm (14, 21)) and could fairly well be modeled by a sum of two OCP forms (*Figure 3A*): an orange OCP^O^ spectrum (~75%) and a violet spectrum with maximum absorption at 550 nm (25 %). Recently, we have demonstrated that the latter violet forms of OCP may appear upon carotenoid (canthaxanthin, CAN) transfer into Apo-OCP (43) and represent dimers of OCP, in which a symmetric CAN molecule with keto-oxygens at both β-rings is symmetrically positioned between two CTDs, as e.g. in the dimeric COCP characterized earlier (15). Such dimers may be additionally stabilized by cross-protein interactions of NTEs and CTDs (36). Formation of such violet forms requires intermediates, in which the carotenoid is positioned in the CTD. Therefore, translocation of the carotenoid from the NTD must occur. We assume that only CAN-coordinating species could be stabilized between two CTDs due to the required symmetry of keto/H-bond interactions. These facts strongly indicate that positioning of the carotenoid in OCP^R^ is not static, and it can move or slide until specific H-bonds with Trp-288 and Tyr-201 are restored, probably, after multiple trials. Due to the absence of the H-bond donor Trp-288, the OCP^OI^ state of the OCPW288A mutant is less stable, but this dynamic equilibrium between OCP^OI^ and OCP^R^ could be shifted towards the orange form by FRP or phosphate, and most likely kosmotropes in general. In the presence of such stabilizing agents, one H-bond with Tyr-201 is sufficient to preserve OCP in the orange-like state. However, neither FRP nor phosphate affect the absorption spectrum of OCP^Y201A/W288A^ mutant, which is similar to such of OCP^W288A^, indicating that in both mutants the carotenoids are not connected to the CTD. It cannot be concluded that the structure of such a phosphate-induced orange-like OCP^W288A^ is identical to OCP^O^, but surprisingly, phosphate does not only restore several of its spectral characteristics, but also the ability to photoconvert under actinic light (*Figure 3B*). This fact is also supported by Raman spectra (*Figure 3C*), which simultaneously exhibit typical features of violet and red OCP forms, but not of the orange state, because it is inevitably photoactivated by 532-nm laser used for Raman spectroscopy. Amazingly, the amplitude of photoconversion in this special case is not reduced by the high phosphate concentration (*Figure 3D*). Rather, the rates of the red to orange back-conversion were inversely proportional to the phosphate concentration (*Supplementary Figure 3*). These facts demonstrate that Trp-288 is not necessary for the photoswitching of OCP itself, but rather is important for the stabilization of the OCP^OI^ intermediate during back-conversion, or for formation of the final compact OCP^O^ state with its distinct carotenoid curvature as recently suggested by Bandara et al. (36).

**Figure 3.**
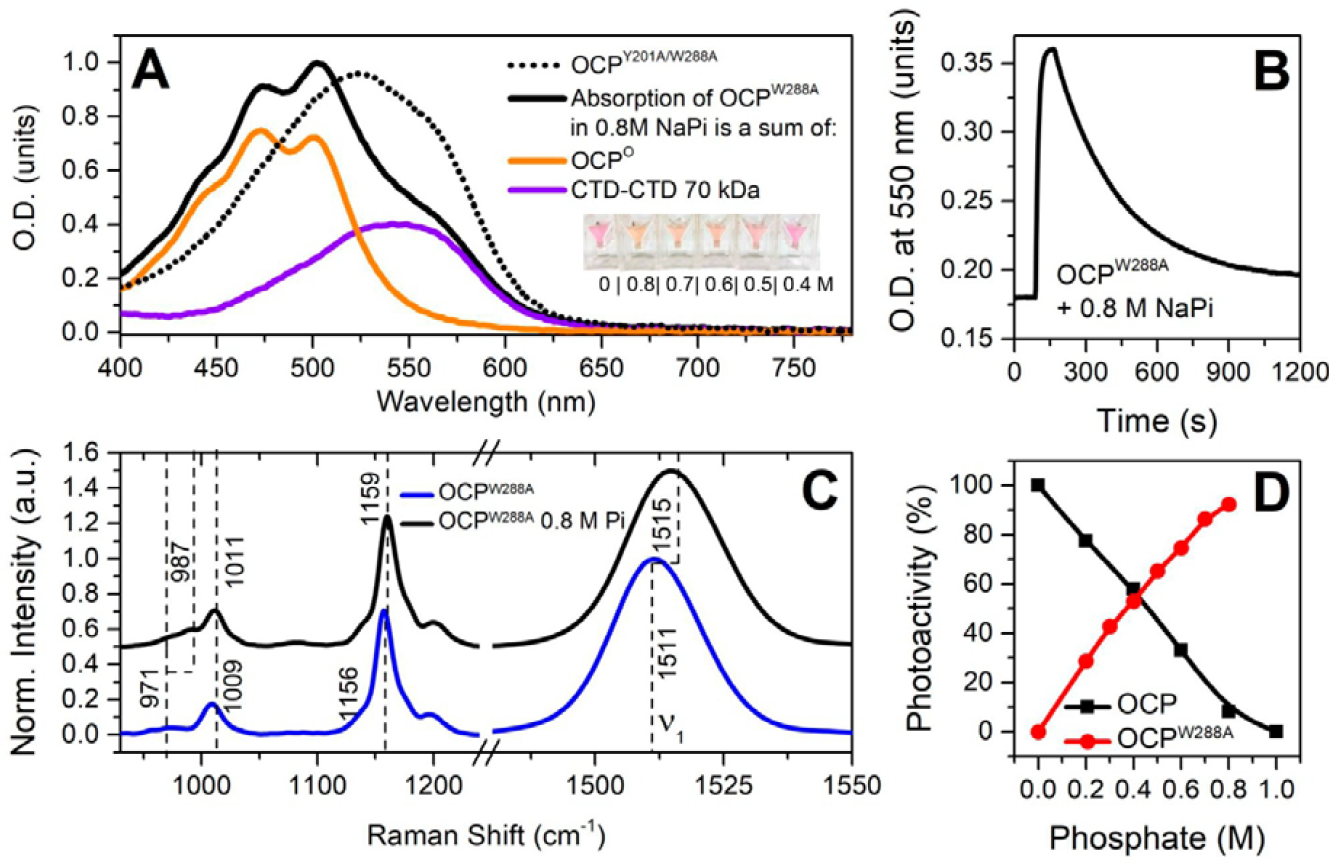
Effects of phosphate on OCP^W288A^. (**A**) - absorption spectrum of OCP^W288A^ in 0.8 M phosphate buffer. The spectrum represents a mixture of orange and violet forms. Dotted line shows absorption of the OCP^Y201A/W288A^ mutant, which identical to the initially purple OCP^W288A^ but not sensitive to phosphate. The inset shows images of cuvettes containing solutions of OCP^W288A^ in different phosphate buffers. (**B**) – photocycle of OCP^W288A^ measured as changes of O.D. at 550 upon illumination by actinic light. (**C**) – Raman spectra of OCP^W288A^ without and with phosphate. (**D**) – photoactivity of the wild-type OCP and OCP^W288A^ mutant at different concentrations of phosphate. Photoactivity was determined as changes of O.D. at 550 nm normalized to concentration of carotenoid in the sample. Maximum changes of O.D. at 550 nm for WT OCP in the absence of phosphate were set to 100 %.

### 2.4 Assignment of tryptophan residues involved in static and dynamic quenching in OCP

The presence of multiple tryptophan residues in OCP (see *Figure 4A*) makes discrimination of their fluorescence properties complicated. This issue cannot be directly solved by mutagenesis due to the significance of Trp residues (in particular, Trp-288, −110 and −41) for functional activity of OCP (18, 19). It is known that excited-state electron transfer to protein backbone carbonyls is the most important mechanism of Trp fluorescence quenching (44). Considering this fact and the tight interactions between Trp-288 and the keto oxygen of the β-ring of ECN, we assume that the fluorescence of Trp-288 could be statically quenched (see above). In our previous work, we have demonstrated that two CTDs of OCP can bind CAN (with keto oxygens at each β-ring) and that Trp-288 plays an important role in stabilization of this homodimeric COCP (15) (43). Comparison of the Trp fluorescence decay kinetics of COCP and its apoprotein form (Apo-COCP), both having only two tryptophans (Trp-277 and Trp-288 in the CTD of *Synechocystis* OCP) reveals that the fluorescence of both Trp residues is significantly quenched by about 85% in the presence of the carotenoid (*Figure 4B*). Trp fluorescence quenching in COCP is of mixed type, with about 25 % dynamic (attributable to the change in average lifetime) and the other about 60 % due to static quenching, which means that in the carotenoid-coordinating homodimer of CTDs, two Trps are statically quenched and the other two contribute to excitation energy transfer (EET) to the carotenoid. We assume that Trp-288 is statically quenched due to H-bonding with the carotenoid and Trp-277 is quenched due to EET. The situation for Trp-277 in COCP might well be different from the one in the OCP^O^ state, because in the latter, Trp-277 could be involved in static quenching, since it forms an H-bond to Asn-104 according to the crystal structure (18), but there is no evidence yet for a similar H-bond in the COCP dimer of CTDs.

For further assignment of the mechanism of Trp fluorescence quenching in OCP, we studied Trp fluorescence in the presence of different concentrations of iodide, a well-known and commonly used collisional (dynamic) quencher. Our idea was to compare the iodide-induced quenching mechanism in different protein forms in order to deduce information about the accessibility of different Trp residues in various OCP states. We found that iodide differently affected the fluorescence decay of OCP in its orange and red forms (*Figure 4C* and *D*). In general, the Stern-Volmer plots (F_0_/F versus iodide concentration) for Apo-COCP, Apo-OCP and OCP^R^ were linear up to 500 mM iodide, with the largest slope observed for Apo-COCP (quenching constant of about 4.5 M^-1^). Essentially coinciding curves with smaller slopes (Stern-Volmer constants of about 3.5 M^-1^) were determined for Apo-OCP and OCP^R^ (see *Supplementary Figure 4*). However, the Stern-Volmer plot of OCP^O^ showed only very limited quenching by iodide with saturating behavior at low iodide concentrations (*Supplementary Figure 4*). This observation suggests a substantially limited accessibility of Trp residues in OCP^O^, which invoked the necessity to evaluate the data according to the modified Stern-Volmer plot (F_0_/(F_0_-F) versus 1/[I^-^], see (45)), as depicted in *Figure 4F*, to determine the fraction of accessible Trp residues (*f_a_*) in OCP^O^. For OCP^O^, the analysis of the modified Stern-Volmer plots yielded a *f_a_* value of about 0.33 (*Figure 4F*). It is important to recall that in the compact OCP^O^ state, three out of five Trp residues are already involved in static quenching due to interactions with the carotenoid (see text above), leaving only two Trp residues contributing to the observed fluorescence amplitude. Only one of these two Trp residues is apparently accessible to collisional quenching by iodide, whereas the other is involved in EET to the carotenoid but without quencher access. Considering the structure of OCP^O^ (PDB entry: 3MG1), the residue accessible to iodide quenching should be Trp-277, which we assume to be involved in EET to the carotenoid together with Trp-101, which is the furthest from the carotenoid, while the Trp residues in the carotenoid-coordinating cavity, Trp-288, Trp-41 and Trp-110, are involved in interactions with the carotenoid leading to the effect of static quenching. This is also in agreement with the character of Trp-277 quenching observed in COCP.

The higher efficiency of Trp quenching in OCP^R^ compared to OCP^O^, as well as for Apo-OCP, which is similarly extended in structure (41), and Apo-COCP, is probably due to differences in protein conformation. While OCP^O^ is highly compact, the red and the other extended states are characterized by domain separation, solvent exposure of the interdomain cavity, and properties of the molten globule state (11, 14), which all increase the accessibility of tryptophans to iodide.

**Figure 4.**
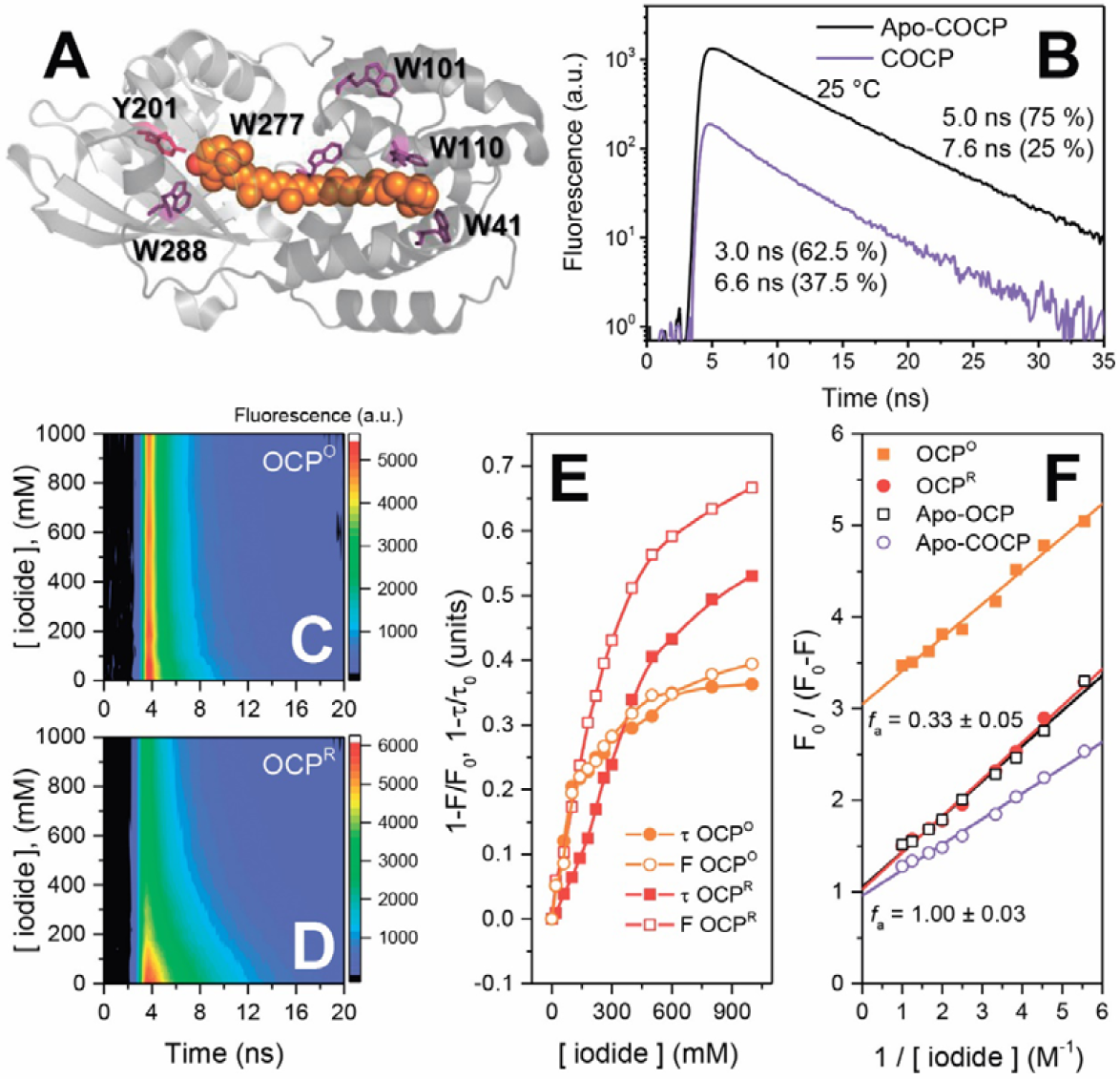
Quenching of Trp fluorescence in OCP and COCP by carotenoid and iodide. (**A**) – location of all Trp and Tyr-201 in the crystal structure of OCP in the orange state (3MG1). Carotenoid is shown by orange spheres, Trp and Tyr-201 residues are shown as violet and pink sticks. (**B**) – fluorescence decay kinetics of COCP (violet) and Apo-COCP (black) measured at 25 °C. Protein concentrations were equal in both samples. (**C**, **D**) – quenching of OCP^O^ and OCP^R^ Trp fluorescence by increasing concentrations of iodide. At each iodide concentration, picosecond fluorescence decay was measured. All experiments with iodide were carried out at 2 °C in order to reduce the rate of OCP^R^ to OCP^O^ back-conversion. (**E**) – dependencies of quenching efficiency on iodide concentration calculated as changes of fluorescence intensities and average lifetimes from the data presented in panels C and D. (**F**) – modified Stern-Volmer plots of Trp fluorescence quenching by iodide approximated by linear functions, the ordinate intercepts define the reciprocal values of the fraction of accessible Trp residues (*f_a_*), as indicated Protein concentrations were equal in all samples.

## 3. Conclusions

In this work, we utilized a combination of time-resolved spectroscopic techniques to study the photoconversion and consequent back-relaxation of OCP. This approach allowed us to determine for the first time two novel intermediate states of the OCP photocycle, which can clearly be identified by distinct spectroscopic features, in addition to the stable orange OCP^O^ and the physiologically active red OCP^R^ (*Figure 5A*). We found that accumulation of the red intermediate state OCP^RI^, which is identical to OCP^R^ in terms of its absorption spectrum, occurs faster than 200 ns. We assume that the extremely fast photoisomerization of the carotenoid disrupts hydrogen bonds between the keto oxygen at the β-ring of ECN (or CAN) and Trp-288/Tyr-201 in the CTD of OCP, which affects the absorption of carotenoid and leads to the first spectral intermediate. Features of Trp fluorescence of this state need to be studied in more detail, but most probably it is already characterized by increased fluorescence intensity due to at least partial elimination of static quenching. The OCP^RI^ state is probably characterized by a protein conformation similar to the dark-adapted OCP^O^, and thus is able to rapidly convert into the orange state by “clamping” the chromophore. Conversion into the red OCP^RI^ form is associated with a (fast) relaxation of the initially bent carotenoid, which, acting just as a spring, may push the protein towards conformational rearrangements including detachment of the αA helix of NTE (36). This reduces the stability of the CTD-NTD protein-protein interactions and leads to separation of domains. Considering the results of dynamic X-ray scattering and atomic structures of OCP^O^ and RCP (26, 36, 46), we may assume that the NTE is acting like a conductor of the signal from the CTD into the NTD, which may trigger small but important rearrangements in the NTD, including the 6° rotation of the α-helix containing Trp-41, and thus facilitating the translocation of carotenoid. In terms of kinetics, the red OCP^R^ state with separated domains is much more stable compared to the intermediate OCP^RI^, since the geometry of OCP^R^ reduces the probability that the carotenoid reaches the CTD again at least 10 000-fold (compare time constants for the OCP^RI^-OCP^O^ (300 μs) and the OCP^R^-OCP^O^ transition (3.3 s)).

Together with the observed kinetic properties of dynamic and static quenching upon back-conversion of OCP^R^ to OCP^O^ (*Figure 2E*), the determination of Trp residues involved in different types of quenching provide novel insights into carotenoid–protein interactions in the intermediate states. Considering the differences in the rates, we postulate that restoration of the initial orange state OCP^O^ occurs in several steps. First of all, the carotenoid restores its position in the NTD, which probably requires a translocation comparable to 12 Å (26). This step does not necessarily affect absorption of the carotenoid-protein system; however, it results in restoration of static quenching of Trp-41 and Trp-110 (*Figure 2E*), and is most probably reversible. We assume that the formation of violet states (*Figure 3A*) is indicative of significant mobility of the carotenoid in the NTD, which can shift by about 12 Å to its position like in the RCP crystal structure (PDB: 4XB4), but it also can slide in the opposite direction. The second step occurs only if NTD and CTD are properly oriented against each other (likely to be assisted by FRP), so that the carotenoid may reach Tyr-201 and Trp-288 in correct configuration, resulting in the formation of the intermediate state OCP^OI^, which is characterized by orange-shifted absorption and static quenching of three Trp residues. This state can be reached without the assistance of Trp-288 (see *Figure 3*), however the latter determines the stability of this intermediate. The next and final step towards OCP^O^ requires adjustments of protein structure which result in restoration of initial efficiency of EET from Trp-277 and Trp-101 to the carotenoid (*Figure 5A*), and most likely also involve the final bending of the carotenoid into its strained configuration as seen in the OCP^O^ crystal structure. These final atomic-scale adjustments of the protein structure probably affect some spectral characteristics of the carotenoid including Raman bands responsible for out-of-plane distortions of the polyene chain due to final compaction of the protein. Thus, a combination of rapid time-resolved spectroscopies, and of infrared spectroscopy in particular, will be indispensable to study such transient species.

Our experiments with the OCP-W288A protein revealed that the carotenoid position in the OCP^R^ state is not confined to only the NTD, and that translocation of the carotenoid from the NTD to CTD may occur spontaneously (see *Figure 3*). Conversion into the orange intermediate OCP^OI^ state is initiated by carotenoid translocation towards the CTD, which probably requires restoration of carotenoid β-ring position and hydrophobic interactions with Trp-41 and Trp-110, simultaneously quenching their fluorescence (*Figure 2E*). Conversion to the orange absorption spectrum is possible due to restoration of the hydrogen bond to Tyr-201 and can be externally stabilized by FRP or high phosphate concentrations, although natural stabilization of carotenoid in this state requires additional bonding with Trp-288. Stabilization of the OCP^OI^ state induces conformational changes of the protein structure, which follows the changes in absorption spectrum at a slower rate, resulting in the pronounced *asynchrony* of the photoactivation-induced changes in OCP (*Figure 2E*). Thus, it is important to discriminate between spectral characteristics and functional (physiological) properties of different OCP states, the changes of which may occur asynchronously. Considering the well-established facts that the carotenoid is an excellent quencher of excitation energy in both red and orange state, and the importance of the OCP conformation for interactions with other proteins, one can hypothesize that OCP^OI^ state is still capable of PBs fluorescence quenching and interaction with FRP, while in terms of its spectrum, this state already largely resembles the initial orange state. We also predict that some OCP mutants, which could be trapped in the orange state, may bind to the phycobilisomes and, probably, even induce quenching. Investigation of such mutants by a combination of time-resolved methods is mandatory to reveal the structural determinants underlying the photoprotection mechanism of OCP-like proteins in general.

**Figure 5.**
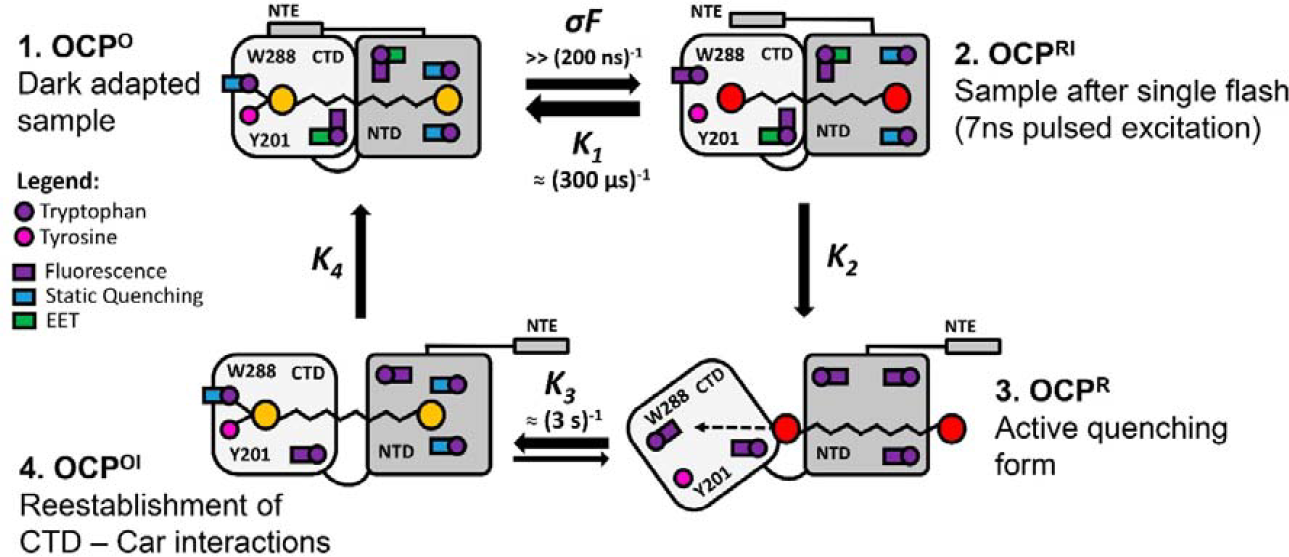
Novel aspects of the OCP photocycle. Absorption of a green photon by the dark-adapted OCP^O^ (state 1) causes excitation and isomerization of the carotenoid molecule, which results in disruption of hydrogen bonds between the keto-group at the β-ring of the carotenoid and Trp-288 and Tyr-201, both located in the CTD of OCP (see *Figure 4*). The immediately appearing red intermediate OCP^RI^ (state 2) is unstable and may rapidly (*K_1_* ~ 300 μs^-1^, see *Figure 1*) return in to the initial orange state, as the protein structure of this red intermediate is similar to OCP^O^. However, carotenoid isomerization induces significant changes of the protein structure, which result in a significantly lower rate of red to orange transition, due to separation of structural domains in OCP^R^ (stage 3). This relatively long-lived red state is physiologically active and can induce quenching of PBs fluorescence. Continuous illumination by actinic light causes accumulation of OCP^R^. However, even under continuous actinic illumination, dissipation of the red form occurs due to the following steps, which include carotenoid translocation into the CTD. This process occurs spontaneously most likely due to a high mobility of carotenoid in its hydrophobic cavity. In the wild-type OCP, this translocation may result in the reestablishment of hydrophobic interactions between one β-ring of the carotenoid and Trp-41 and Trp-101 in the NTD. Moreover, hydrogen bonds between the keto oxygen of the carotenoid and Tyr-201 and Trp-288 are formed, resulting in stabilization of the cofactor in the orange intermediate state OCP^OI^ (stage 4), in which fluorescence of Trp-41, 101 and 208 is statically quenched, but the energy transfer to Trp-110 and Trp-277 is not yet established (see *Figures 2* and *4*). The latter is brought about by further rearrangement of the protein. The location of the carotenoid in the CTD is destabilized in the W288A mutant, so this sample is trapped between the stages 3 and 4, due to the presence of only one hydrogen bond between the carotenoid and Tyr-201. The presence of FRP or high concentration of phosphates increases the probability of intermediate OCP^OI^ state formation. Absence of both Trp-288 and Tyr-201 traps the sample at stage 3.

## 4. Materials and Methods

### 4.1 Protein purification

Cloning, expression and purification of the 6xHis-tagged *Synechocystis* OCP, OCP lacking the NTE comprising the first 20 amino acids (ΔNTE), and FRP were described previously (14, 21, 22). Holoforms of the 6xHis-tagged OCP were expressed in echinenone and canthaxanthin-producing *Escherichia coli* cells essentially as described before (15). The His-tag was identical in all the protein forms used. All proteins were purified by immobilized metal-affinity and size-exclusion chromatography to electrophoretic homogeneity and stored at +4 °C in the presence of sodium azide.

### 4.2. Steady state absorption measurements

Absorption spectra were recorded using a Maya2000 Pro (Ocean Optics, USA) spectrometer as described in (11, 21, 28). Upon absorption measurements, a blue light-emitting diode (LED) (M455L3, Thorlabs, USA) with a maximum emission at 455 nm was used for the photoconversion of the samples (further referred to as actinic light for OCP^O^→OCP^R^ photoconversion). The temperature of the sample was stabilized by a Peltier-controlled cuvette holder Qpod 2e (Quantum Northwest, USA) with a magnetic stirrer.

### 4.3. Measurements of flash-induced absorption transients

Flash-induced absorption changes were examined with a lab-made flash-photolysis system. Photocycle excitation was provided by the second harmonic of a Nd–YAG Q-switched laser LS-2131M (LOTIS TII, Belarus), using 7-ns pulses at 532 nm. A monochromatic beam with 550 nm and 6 nm FWHM, which was obtained with two monochromators (before and after the sample) from a power-stabilized 100 W halogen lamp (OSRAM, Germany), was used for monitoring transient absorption.

Absorption changes were followed by using a 9658 B (Electron Tubes Ltd., UK) photomultiplier with 300-MHz wide-bandwidth amplifier and a 100 MHz Octopus CompuScope 8327 (Gage, Canada) analog-to-digital converter. Five hundred single traces were averaged to achieve a higher signal-to-noise ratio. The duration between flashes was set to be in the range between 1 and 5 s to allow completion of a photocycle. The traces were converted into a quasilogarithmic time scale by using home-made software. The kinetic traces were fit with a sum of exponentials using Mathematica (Wolfram Research, USA) or OriginPro 2015 (Originlab, USA).

### 4.4. Time Correlated Single-Photon Counting (TCSPC) techniques for tryptophan fluorescence transients

In all experiments, OCP photoconversion was triggered by a blue LED (M455L3, Thorlabs, USA), which passed through the FB450-40 bandpass filter (Thorlabs, USA). Fluorescence of the samples was excited by picosecond EPLED-280 laser diode (Edinburgh Instruments, Scotland) with maximum emission at 280 nm corrected by a bandpass filter (AHF, Germany), delivering pulses at 10 MHz repetition rate. Fluorescence of samples in the range from 300 to 500 nm was collected at 90° to excitation pulse via a collimation lens coupled to the optical fiber of the 16-channel detector PML-16-C (Becker&Hickl, Germany). Fluorescence was recorded with a time- and wavelength-correlated single photon counting system SPC-150 (Becker&Hickl, Germany) described in (28). Thus, series of fluorescence decay kinetics were obtained during different stages of back-conversion of the samples after exposition of the sample to actinic light. The temperature of the sample was stabilized by a Peltier-controlled cuvette holder Qpod 2e (Quantum Northwest, USA) with a magnetic stirrer.

The fluorescence decay curves were approximated by a sum of exponential decay functions with the SPCImage (Becker and Hickl, Germany) software package. To compare different decay curves, we calculated the average decay time according to the expression: 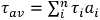, where *τ_i_* and *a_i_* are the lifetime and the amplitude (normalized to unity: 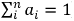) of the *i*-th fluorescence decay component. To obtain the time integrated fluorescence intensity, the number of photons in each time channel of kinetics was summed up. All calculations were performed using the Origin 9.1 (OriginLab Corporation, USA).

For steady-state fluorescence emission and fluorescence excitation spectra measurements, a FluoroMax-4 instrument (Horiba Jobin Yvon, Japan-France) was used.

All experiments presented in this work were repeated at least five times.

## Acknowledgements

This work was supported by grants from Russian Foundation for Basic Research 15-04-01930a to E.G.M. and 15-29-01167 to V.Z.P. E.G.M. was supported by Dynasty Foundation Fellowship. The reported study was funded by RFBR and Moscow city Government according to the research project № 15-34-70007 «mol_ а_mos». T.F. acknowledges the support by the German Federal Ministry of Education and Research (WTZ-RUS grant 01DJ15007) and the German Research Foundation (Cluster of Excellence “Unifying Concepts in Catalysis”).

## Supplementary Information

**Figure S1.**
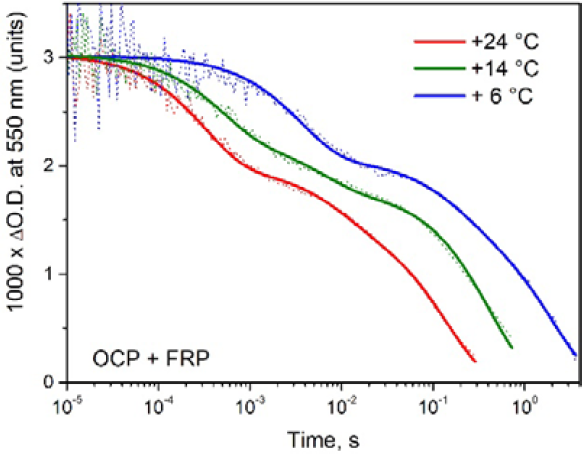
Characteristic flash-induced transitions of WT OCP absorption at 550 nm at different temperatures approximated by multiexponential decay. All samples contained OCP and FRP (1/1.6 concentration ratio). The absence or low efficiency of excitation energy transfer (EET) in the red state could be at least partially explained by the changes of overlap integral between the Trp emission (donor) and ECN absorption (acceptor), which decreases due to the red shift of ECN absorption in the photoactivated state. Another reason could be the increased mobility of carotenoid molecule in the red state, which may lead to reduction of the role of (specific) orientation factors between Trp residues and the carotenoid. It should be noted that at least at low temperatures, the average Trp fluorescence lifetimes of OCP^R^ are significantly larger compared to that of the OCP apoprotein (Apo-OCP) (*Supplementary Figure 2*), indicating that in this state quenching is not only removed, but that the carotenoid provides a hydrophobic environment for some Trp residues, which would shield them from collisional interactions with water compared to the situation in Apo-OCP. Previously it was shown that the largest solvent accessibility changes after photoconversion occurred in CAN binding residues, namely a decrease in solvent accessibility of Trp-41 (26).

**Figure S2.**
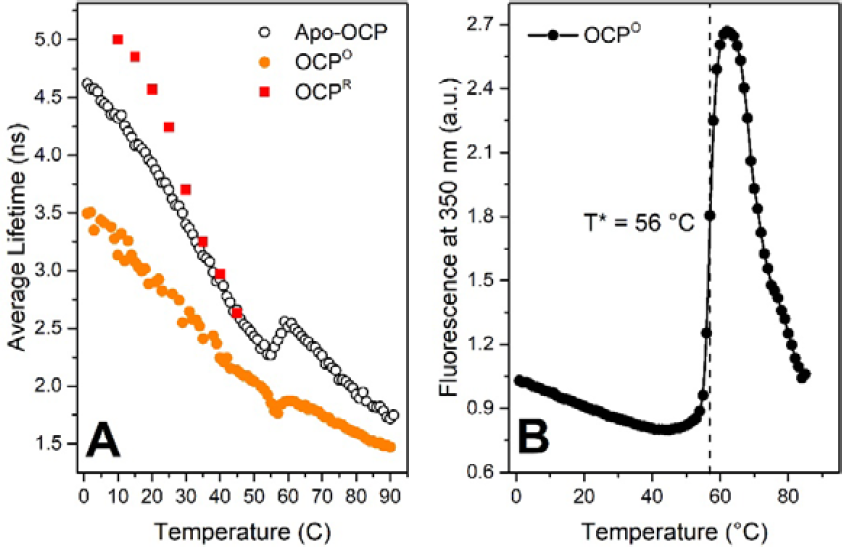
(**A**) - temperature dependency of tryptophan average fluorescence lifetime of OCP^O^, OCP^R^ and Apo-OCP registered at 350 nm. (**B**) – temperature dependency of fluorescence intensity of OCP^O^ T* indicates half-transition (melting) temperature.

**Figure S3.**
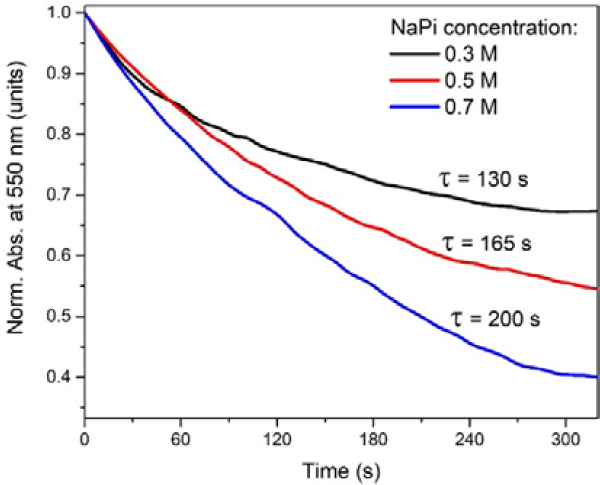
Time-courses of red to orange back-conversion recorded as absorption changes at 550 nm of initially non-photoactive purple OCP-W288A mutant in presence of different concentrations of phosphate. The Stern-Volmer plots of Apo-OCP and OCP^R^ also exhibited some slight negative deviation from linearity at iodide concentrations above 500 mM, which on the first sight also suggests limited accessibility of Trp residues; however, the modified Stern-Volmer plots for these species (*Figure 4F*) yield an accessible fraction (*f_a_*) of 1. This indicates that some other effect limits the quenching at high concentrations of iodide. Since iodide anions belong to the group of chaotropes in the Hofmeister series, it is possible that high iodide concentrations induce slight (local) destabilizations of the protein structure, which could entail changes in the microenvironment of one or the other Trp residue.

**Figure S4.**
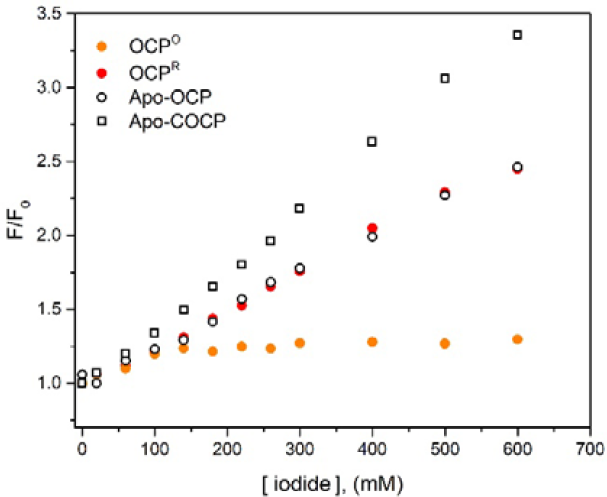
Stern-Volmer plots for Apo-COCP, Apo-OCP, OCP^O^ and OCP^R^.

